# Experimental reduction of land use increases invertebrate abundance but not diversity in grasslands

**DOI:** 10.1101/2025.03.14.643238

**Authors:** Michael Staab, Alexander Keller, Rafael Achury, Andrea Hilpert, Norbert Hölzel, Daniel Prati, Wolfgang W. Weisser, Nico Blüthgen

## Abstract

Grasslands are diverse ecosystems that are increasingly threatened by intensive land use. Restoring grasslands by reducing land-use intensity may support insect abundance and diversity, helping to halt insect declines. To test for the effect of reduced land use on invertebrates, we studied an experiment (established 2020) at 45 sites across three regions of Germany. We hypothesized that reduced land use increases invertebrate abundance and diversity, with larger effects in less intensively used grasslands. Using suction sampling, invertebrates were quantitatively sampled in May 2021 and May 2023, with 2021 samples identified by DNA meta-barcoding. Reducing land use to a single late mowing increased invertebrate abundance by 41% after one year and 99% after three years. However, species richness, Shannon diversity, and Simpson diversity did not differ between treatments and controls. Finding more individuals in grasslands with reduced land use suggests that species already present benefit, rather than additional species being recruited from the surrounding area. The effect of land-use reduction on abundance was consistently influenced by land use in the surrounding matrix, with larger positive effect sizes at grasslands with lower mowing frequency but higher fertilization. In spite of these local differences in the magnitude of restoration effects, the consistent increase in invertebrate abundance suggests that reducing land-use intensity can enhance invertebrate populations with potential benefits for ecosystem functions. It will be important to study how outcomes of land-use reduction develop over time, as land-use reduction is likely more successful when implemented permanently.

## Introduction

Habitat loss and ecosystem degradation driven by intensive land use, particularly agriculture, are threats to biodiversity globally (Outhwaite et al., 2022). The intensive farming practices in agricultural land alter habitat structure, disrupt ecological processes and results in large-scale declines of many species (Butchart et al., 2010). A particularly alarming consequence is the widespread decline in insects and other invertebrates (Seibold et al., 2019; Staab et al., 2023; Wagner, 2020), which are vital for many processes and functions in ecosystems (Weisser & Siemann, 2004). In addition to preserve the remaining natural areas (Watson et al., 2016), restoring habitats to halt and eventually reverse ecosystem degradation and species decline is one of the great ecological and political challenges of the 21^st^ century, reflected e.g. by the concurrent UN Decade on Ecosystem Restoration (United Nations Environment Programme, 2021).

Grasslands, which support a rich diversity of plant and animal species, are especially vulnerable to land-use intensification and hence in need of restoration (Lyons et al., 2023). Current grassland management practices frequently maximize productivity at the expense of biodiversity and other ecosystem functions (Gossner et al., 2016). Land use in, both, natural and semi-natural grasslands consists of a combination of fertilization, mowing and grazing, which when applied with high intensity, favor productive but species poor and compositionally less diverse plant communities (Busch et al., 2019; Hautier et al., 2009). The outcomes of these different land-use components may negatively influence species separately but also jointly (Chisté et al., 2016; Wagner et al., 2021). For example, each mowing event imposes high mortality for arthropods (Berger et al., 2024b; Humbert et al., 2009; Steidle et al., 2022) while fertilization reduces plant diversity via increased productivity and competition for light, effects on arthropods are rather indirect (Nessel et al., 2023; Prestidge, 1982). Nevertheless, frequency of mowing and intensity of fertilization are often inherently correlated, as more productive fertilized grasslands are cut more often (Blüthgen et al., 2012; Vogt et al., 2019). Similarly, mowing and grazing both reduce plant biomass and hence remove resources and shelter for insects (Berger et al., 2024b; Helden et al., 2020; Kruess & Tscharntke, 2002), but effects of frequent mowing are considerably more detrimental than of grazing, except at very high stocking with livestock (Fartmann, 2024).

Restoring intensive towards more extensive grasslands by reducing land-use intensity can preserve and increase insect abundance and diversity (Fartmann, 2024; Gossner et al., 2016; Kruess & Tscharntke, 2002). For plants a key strategy to restore diversity when nutrient availability is high involves limiting nutrient inputs and removing biomass to lower the competitive ability of dominant plant species (Andraczek et al., 2023; Eskelinen et al., 2022; Hautier et al., 2009). However, it cannot be assumed that grassland management aiming at promoting plant diversity will be equally beneficial for invertebrates. Biomass removal, especially by mowing, will be negative for insects because every single mowing event exerts substantial mortality risk (Berger et al., 2024b; Humbert et al., 2009). Most past studies investigating relationships between land-use intensity and insect abundance or diversity were observational or correlational (Allan et al., 2014; Frenzel & Fischer, 2022; Gossner et al., 2016). Yet, to rigorously establish causality and to disentangle mechanisms larger-scale field experiments are needed (Weisser et al., 2023). Very few regionally-replicated experiments have studied the reduction of land-use intensity in grasslands, and usually focused on plants (Humann-Guilleminot et al., 2023; Van Vooren et al., 2018). Nevertheless, to account for potential local context dependencies, such as differences in management or landscape configuration, experiments that compare the effects of land-use reduction on invertebrates to spatially proximate control with regular land-use intensity are desirable. These studies will help understand if and how grasslands can be managed to restore insect populations after the declines seen in recent decades.

To test for the effect of reduced land-use intensity on invertebrates we utilized a recently established extensification experiment (Andraczek et al., 2023) that has been implemented within the framework of the Biodiversity Exploratories across three regions of Germany (Fischer et al., 2010). We hypothesized that experimentally reduced land use increases invertebrate abundance and diversity (Fig. 1), with the magnitude of the increase (in regard to control) being larger at formerly less-intensively used grasslands.

**Fig. 1.**
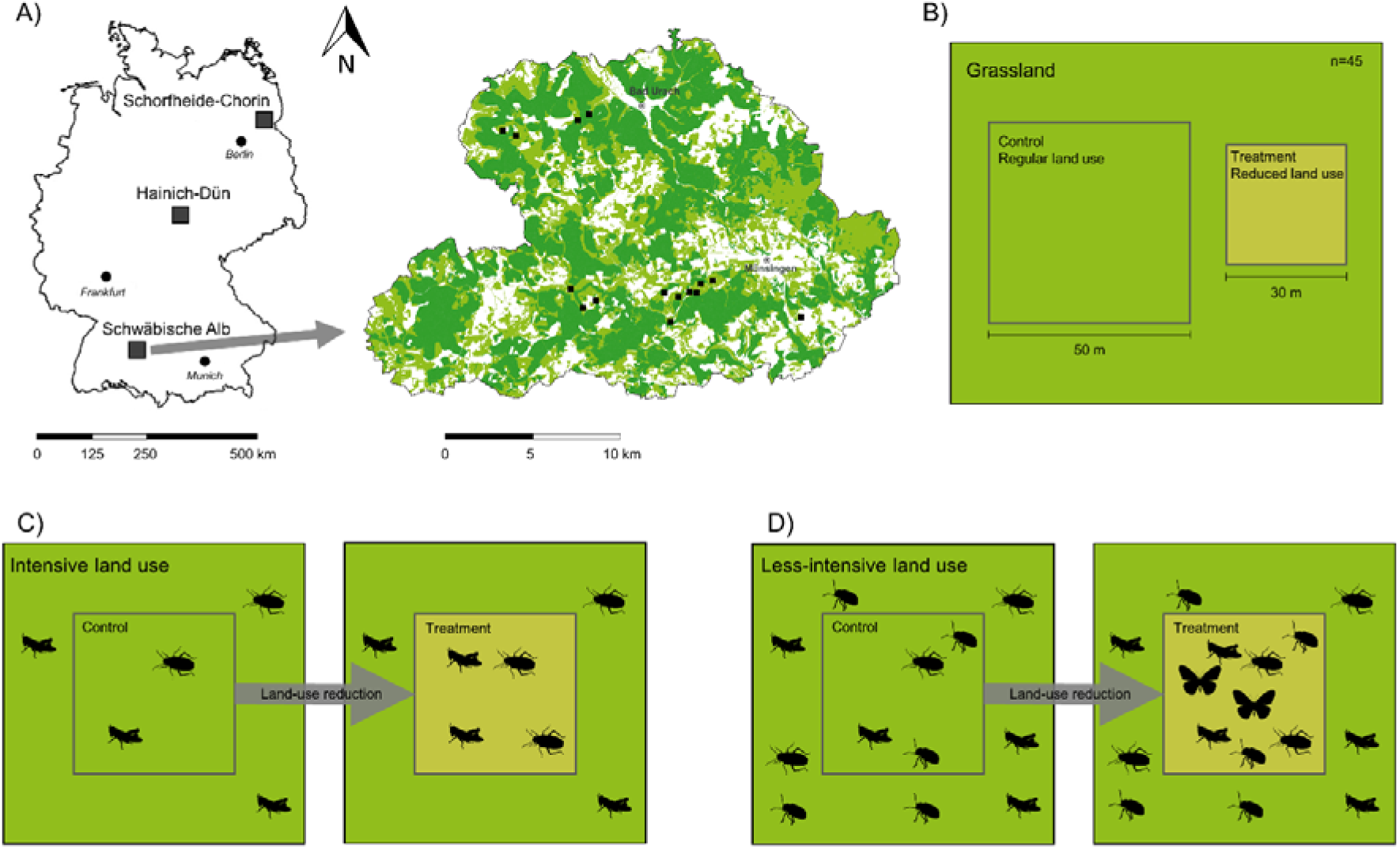
Overview on study design and hypothesis. A) Across three regions of Germany, land-use reduction experiments were established. Depicted for the Schwäbische Alb region, black squares show location of experiments, light green indicates grasslands, and dark green forests (with white being agriculture and settlements). B) Per region, land use at 15 grasslands each (total n=45) was experimentally reduced within a matrix that also comprised the control plot. C, D) We expected that the experimental reduction of land use increases the abundance and diversity of arthropods and that the increase is larger in less-intensively used grasslands. Maps are from OpenStreetMap (OpenStreetMap Contributors 2022) and arthropod silhouettes from phylopic.org (available under a CC0 1.0 license).

## Materials and methods

### Study sites

The study was conducted in the tree German regions Schwäbische Alb, Hainich-Dün and Schorfheide-Chorin (Fig. 1) within the Biodiversity Exploratories consortium (Fischer et al., 2010). The regions differ in climate and topography, making them representative for a large part of Central Europe. In each region, 50 grassland study sites (50 m x 50 m) were established in 2008 and have since then been monitored for land-use intensity and arthropod communities, among many other aspects of biodiversity and ecosystem functioning (www. https://www.biodiversity-exploratories.de). Local grasslands are farmed as meadows (mown at least once, if grazed then only briefly by sheep), pastures (only grazed, not mown yearly) or mixed pastures (mowing and grazing in the same year), with no influence of researchers on any decision concerning land use. All study sites are embedded in larger management units and comprise a gradient of land-use intensity typical of each region, from unfertilized extensive pastures to intensively managed silage grasslands. Land-use intensity (Blüthgen et al., 2012) is assessed yearly via standardized questionnaires with farmers (Vogt et al., 2019), from which the frequency of mowing (yearly number of cuts), intensity of grazing (livestock units per ha and year) and fertilization (kg N per ha and year) are calculated. Further details on land use (e.g. exact mowing dates, mowing machinery) are also recorded. The size of each grassland (in ha), which represents a management unit receiving the same land use, was measured from land property maps. The proportion of grassland and agriculture within a 500 m radius around each plot was retrieved from satellite-based vectorized ATKIs Basis DLM land cover data (Seibold et al., 2019).

### Land-use reduction experiment

To investigate how land-use reduction relates to biodiversity and ecological processes, a controlled experiment was implemented at 15 sites per region (total n=45) before the start of the vegetation period in 2020 (Andraczek et al., 2023; Weisser et al., 2023). For this, sites with medium to high land-use intensity were selected, which could be experimentally reduced. In spatial proximity of the long-term monitoring plot used as control with regular land use, a plot (size 30 m x 30 m) with experimentally-reduced land-use was established (Fig. 1). This reduced land-use plot was subsequently only mown once per year late in the growing season in August or September, without further land use such as fertilization, grazing or more mowing. Information on mowing dates and mowing machinery (e.g. cutting height in cm; use of a conditioner, which further crushes grass after being cut) was noted. All experimental plots were mown with rotary mowers, but the technical details such as cutting height and the date of mowing varied among plots. Further details on the experiment including plant species composition data, which did not differ between reduced land use and control one year after starting the experiment, are available in Andraczek et al. (2023).

### Invertebrates sampling

Invertebrates were collected in the reduced land-use and control plots (Appendix A: Fig. S1) using a custom-built suction sampler modified from a leaf vacuum (Stihl SH 86, STIHL, Waiblingen, Germany). Sampling was done one year after the start of the experiment in May 2021 prior to the first mowing event or the first intensive grazing at all control sites. Sampling before the first mowing or grazing is necessary to ensure that reduced land-use plots are not simply acting as refuges for arthropods within an unfavorable matrix (Berger et al., 2024a; Schwarz et al., 2023). In each plot, two 1 m x 1m subplots (5 m apart) were predetermined near the center of the plot. For each sampled subplot, vegetation height was measured at five points with a folding rule and averaged. An aluminum cage (biocenometer) wrapped in fine insect net to prevent the escape of specimens was placed on each subplot, and all invertebrates were sucked from the 1 m² area for five minutes into a mesh bag. Specimens were immobilized with carbon dioxide, and stored at -18°C until further processing. Arthropods (and gastropods, here jointly referred to as invertebrates) were separated from soil and plant debris under a stereomicroscope, counted at least on order level, and preserved in 100% ethanol. In total, 176 samples were taken in 2021 (44 sites, 2 treatments, 2 subplots) as one of the 45 experimental sites could not be sampled. Before analysis and species identification by meta-barcoding, samples from both subplots per treatment plot for each year were combined (Melcher et al., 2024). The exact same sampling at precisely the same subplots was repeated in May 2023, three years after the start of the experiment (all sites sampled).

### Meta-barcoding

To delineate species identities in 2021 bulk samples (2023 could not be meta-barcoded due to budget constraints), DNA meta-barcoding through next generation sequencing was applied (Morinière et al., 2016). Using meta-barcoding has the advantage over classical morphological species identification that larval stages, which for several orders (e.g. Hemiptera, Orthoptera) dominated in the samples due to the necessary early sampling, can be identified. Furthermore, taxa for which taxonomic expertise is scant (e.g. many Diptera) can be included, allowing a comprehensive assessment of the effects of reduced land use on invertebrates. Meta-barcoding was performed by AIM (Leipzig, Germany). Sample homogenization and tissue lysis was done for 8 h at 56°C with a 10:1 mixture of lysis buffer and proteinase k. For DNA extraction, a 500 µL aliquot was used and a standard protocol for high throughput DNA extraction applied (Ivanova et al., 2006). Purified DNA was eluted in 50 µL molecular water. The 5P region of the mitochondrial COI gene was amplified with the primers mlCOIntf and dgHco (Leray et al., 2013) with Illumina adaptors on the 5’end applying the PCR conditions described in Morinière et al. (2016). PCR products were analyzed through gel electrophoresis, cleaned and re-eluted. Concentration of DNA was measured with a fluorometer (Qubit 4, Thermo Fisher Scientific, Waltham, USA). Sequencing was done on an Illumina 2 x 250 bp MiSeq machine (Illumina, San Diego, USA) at approximately 150,000 reads per sample. More details on all laboratory procedures are available in Leroy et al. (2022).

Bioinformatic processing followed version 400d87c of the pipeline from Leonhardt et al. (2022) (available at https://github.com/chiras/metabarcoding_pipeline). The pipeline utilized VSEARCH v2.18.0 (Rognes et al., 2016) and conducted paired-read merging, quality (ee < 1) and length (200-500 bp) filtering and truncation, primer and adapter removal, dereplication and denoising to amplicon sequences variants (ASV; Eren et al., 2013). Compared to traditional operational taxonomic units (OTUs), ASVs are independent of similarity thresholds and less prone to erroneous classifications that may inflate species numbers (Callahan et al., 2017). Two samples had problems with sequencing and these samples were not used in analyses. Taxonomy was assigned to ASVs with an iterative approach, with first direct global alignments and a threshold of 97% against localized databases. These databases were created using BCdatabaser (Keller et al., 2020) with invertebrate lists of Germany and Central Europe. Sequences were classified first against the German reference database, unmatched sequences then against Central European and finally such still unclassified against entire BOLD (Ratnasingham & Hebert, 2007). Remaining unclassified reads were hierarchically classified using SINTAX (Edgar, 2016) against BOLD to get taxonomic levels as deep as possible with a threshold of 0.8.

Of all ASVs, only sequences classified as invertebrate animals (phyla Arthropoda and Mollusca) were retained (i.e. plants and fungi removed). Furthermore, unspecific classifications that were not at least resolved at the level of taxonomic family were excluded. If multiple ASVs were belonging to the same biological species, these were collated per sample. Read numbers were converted to relative data. We applied low-abundance filtering by discarding ASVs that contributed to less than 0.2% of relative reads per sample, which removes sequences that are likely artefacts or contamination. For comparison, low abundance filtering was also performed for 0.1% and 0.5%. Diversity variables for these alternate thresholds for low-abundance filtering were for the respective same variable highly correlated with diversities based on the 0.2% threshold (Appendix A: Fig. S2; Spearman’s ρ > 0.71), and yielded congruent statistical results.

### Data analyses

All analyses were conducted in R 4.3.1 (www.r-project.org) applying the packages ‘car’ (Fox & Weisberg, 2019), ‘emmeans’ (Lenth, 2024), ‘lme4’ (Bates et al., 2015), ‘MuMIn’ (Bartoń, 2024), ‘phyloseq’ (McMurdie & Holmes, 2013) and ‘vegan’ (Oksanen et al., 2022). The effect of the reduced land-use treatment was expressed as log-response ratio log(treatment/control) (Hedges et al., 1999), with positive values indicating a positive effect of reduced land use (e.g. more individuals) compared to the control, while negative values indicate a negative effect. For abundance, log-response ratios were calculated without Acariformes (mites), Collembola (springtails) and Formicidae (ants). Soil mesofauna (mites and springtails) is not assessed representatively with suction sampling. For ants, as colonial organisms, specimen numbers can be highly inflated when an ant nest is present in the sampling area, which is why they are regularly excluded from abundance analyses (see also Steidle et al., 2022).

From the species data identified by meta-barcoding, invertebrate species richness, exponential Shannon diversity, and Simpson diversity were calculated with ‘phyloseq’. These three measures of diversity follow the Hill-series (species richness: q=0, Shannon diversity: q=1, Simpson diversity: q=2), with a decreasing influence of rare species on the respective diversity (Hill, 1973). Due to one site in 2021 not being accessible and two samples with failures in sequencing final sample sizes were 44 (abundance after one year), 42 (diversities and composition) and 45 (abundance after three years). To evaluate sampling efficiency, species accumulation curves (sample based, 10,000 permutations) and first-order jackknife species richness estimators were computed for the full data and separately for the three regions Schwäbische Alb, Hainich and Schorfheide-Chorin (‘vegan’). To test if treatment effects were different from null, univariate linear mixed-effects models treating region as random intercept to account for possible region-specific response to land-use reduction were used (‘lme4’). Likewise, differences in magnitude of treatment effects (after one year) among abundance, species richness, Shannon diversity and Simpson diversity were tested with a linear mixed-effects model (region as random intercept) and subsequent pairwise contrasts that were Bonferroni-Holm adjusted for multiple comparisons (‘emmeans’). With the same framework, differences in treatment effects on abundance between 2021 (one year after establishment of the experiment) and 2023 (three years after) were assessed.

To test if the magnitude of land-use reduction effects depended on land use in the surrounding matrix, the wider landscape, or specific management decisions on the reduction plot, separate linear mixed-effects models (‘lme4’) were calculated for log-response ratios of abundance (after one and after three years), species richness, Shannon diversity and Simpson diversity. Region was treated as random intercept. All models contained as fixed effects the intensities of mowing, fertilization and grazing of the surrounding matrix (including control plot), each averaged over the three years prior to sampling; the size of the continuous grassland unit, and the proportional cover of grassland and arable land (500 m radius) surrounding the sites. Fixed effects to characterize the management of the reduced land-use plot, were the cutting height of the treatment, if a conditioner was used during mowing, and if treatment and control plot, and hence the surrounding matrix, were mown on the same day. Further, the difference in average vegetation height, also expressed as log-response ratio between treatment and control, which may modulate habitat availability for invertebrates, was included. Mowing and grazing intensity were square-root transformed and the size of the grassland unit was log-transformed to increase normality of the data. All numerical fixed effects were scaled and centered (mean=0, SD=±1) to allow direct comparison of statistical effect sizes. Because each of these fixed effects may attenuate the treatment effect and since for each response variable there were multiple candidate models with different combinations of fixed effects that had a comparable fit to the data (i.e. _Δ_AICc<2, Akaike Information Criterion), we employed model averaging (‘MuMIn’). This involved calculating a model for every possible combination of fixed effects, and then averaging all candidate models within 2 AICc units of the model with the lowest AICc value, applying the full average weighted according to each model’s relative explanatory power. Collinearity among the fixed effects was assessed with variance inflation factors (‘car’) that were consistently <3.5 in all averaged models, making collinearity unlikely (Dormann et al. 2013). Model residuals met assumptions of normality and heterogeneity of variances.

Because experimental reduction of land use may not only affect invertebrate abundance and diversity but likewise species composition, differences in composition between treatment and control were calculated as pairwise Bray-Curtis dissimilarities based on normalized relative reads. Pairwise dissimilarities were then analyzed as response variable in a linear mixed-effects model that had the same fixed effects and random effect, and were subjected to an identical model averaging procedure as the models for log-response ratios described before. Variation in species composition was visualized using 2-dimensional non-metric multidimensional scaling (NMDS), based on Bray-Curtis dissimilarities of normalized reads (‘vegan’). To test for differences in multivariate distances between treatment and control, a permutational multivariate analysis of variance (PERMANOVA) was used. Multivariate homogeneity of group (treatment, control) dispersions was assessed with a permutational beta-dispersion test. To examine the relationship between species composition and treatment the scores of the first two NMDS axes were correlated with (all variables were scaled) average vegetation height, continuous grassland size (log-transformed), grassland cover, arable land cover, mean average vegetation height, mowing intensity, grazing intensity (square-root-transformed) and fertilization intensity (square-root-transformed), using a post-hoc permutation test. All permutational procedures used 10,000 permutations that were stratified within each region (analogous to a random effect).

## Results

In total, 83,343 invertebrate individuals were collected in 2021 (after one year) and 115,490 individuals in 2023 (after three years; each excluding ants, mites, springtails). Meta-barcoding for 2021 specimens considering only sequences contributing to more than 0.2% of relative reads per sample identified 479 unique species (based on ASVs). Coleoptera (beetles) and Diptera (flies and midges) were with, respectively, 130 and 124 species most species-rich (Appendix A: Table S1). Total sampling efficiency was 72.3%, with 663 ± 27 SE (first-order jackknife estimator) species expected at infinite sampling. Sampling efficiency did not differ between reduced land-use (68.4%) and control plots (69.5%), and was comparable across the three regions (range: 68.4-72.1%; Appendix A: Fig. S3). Almost all species recorded by meta-barcoding had been found on the sites before (Appendix A: Fig. S4).

On average, experimentally reducing land use to one late mowing increased abundance after one year by 41% from a mean of 786 ± 464 SD individuals to 1108 ± 468 individuals (Table 1). However, species richness, Shannon diversity and Simpson diversity between treatment and control were almost identical (Table 1). After three years, abundances in reduced land-use plots were 99% higher (1708 ± 726 vs. 859 ± 577). Treatment effects, quantified as log-response ratio, after one year were larger than null for abundance (t=4.954, p<0.001) but not for species richness (t=0.975, p=0.335), Shannon diversity (t=0.900, p=0.373) or Simpson diversity (t=1.035, p=0.307) (Fig. 2A). Consequently, the treatment effect for abundance was larger (all Bonferroni-Holm corrected pairwise contrasts p<0.001) compared to species richness, Shannon diversity and Simpson diversity which did not differ (pairwise contrasts among the latter three p=1.000) (Appendix A: Table S2). After three years, the treatment effect on abundance was also positive (t=8.191, p<0.001), which was larger compared to after one year (t=3.698, p<0.001) (Fig. 2B).

**Fig. 2.**
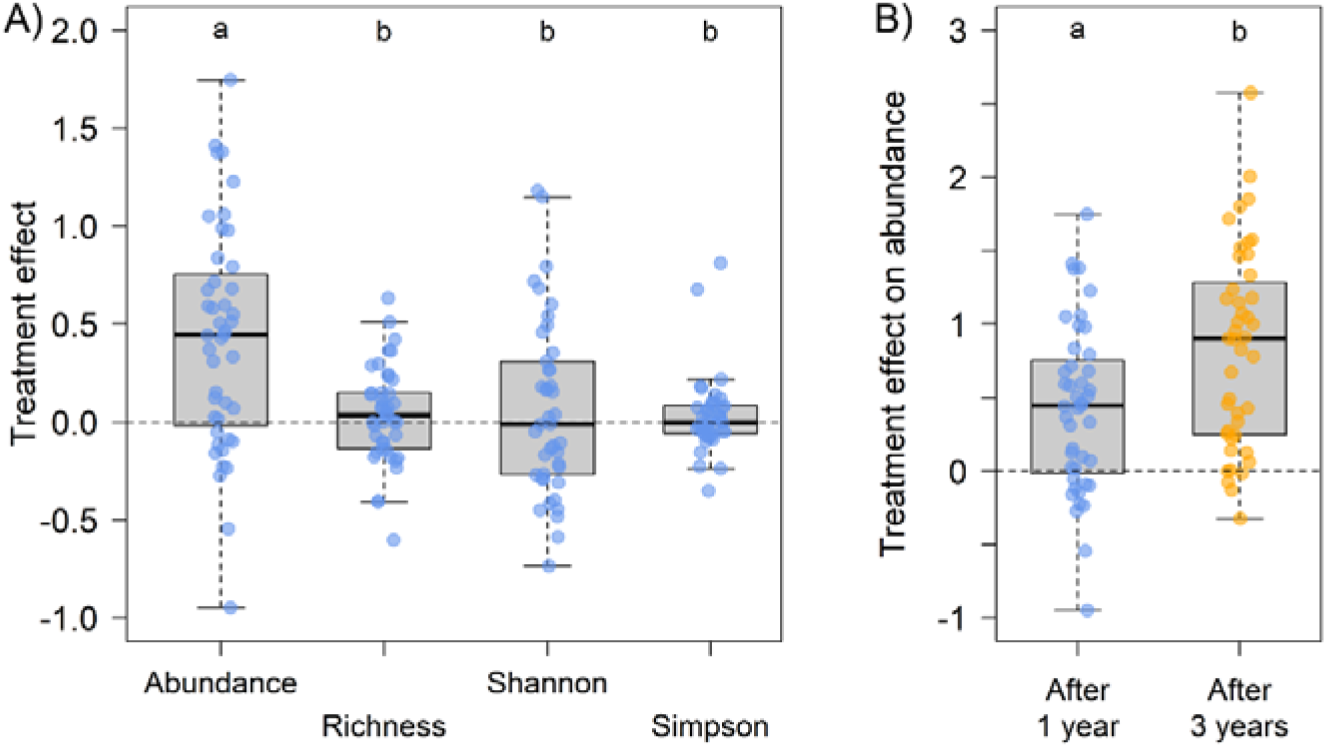
The experimental reduction of land use had A) after one year (blue data) a positive effect (quantified as log-response ratio) on invertebrate abundance but not species richness, Shannon diversity, and Simpson diversity. B) Effects on abundance increased over time, being larger after three years (golden data) than after one year. Values larger than null (dashed horizontal lines) indicate a positive effect of experimental land-use reduction. Abundance data exclude mites, springtails and ants (see methods). Species richness, Shannon diversity and Simpson diversity are based on meta-barcoding reads accounting for more than 0.2% of normalized reads per sample. Lower case letters above boxes indicate significant differences (p<0.001) among treatment effects. Note that the y-axes display different ranges.

**Table 1.** Summary information of the experimental reduction of land use at 45 grassland sites on invertebrate abundance (1 and 3 years after the experiment was established), species richness, Shannon diversity and Simpson diversity (each after 1 year; considering meta-barcoding reads accounting for more than 0.2% of normalized reads per sample). Data always refer to a sampling area of 2 m². Abundance excludes mites, springtails and ants (see methods).

| Variable | Plot type | Range | Median | Mean ± SD |
| --- | --- | --- | --- | --- |
| Abundance after 1 year | Reduced land use | 322 – 2414 | 1029 | 1108 ± 468 |
| | Control | 183 – 2003 | 620 | 786 ± 464 |
| Species richness after 1 year | Reduced land use | 17 – 54 | 32 | 35 ± 9 |
| | Control | 16 – 58 | 33 | 34 ± 11 |
| Shannon diversity after 1 year | Reduced land use | 2.9 – 26.6 | 12.3 | 12.8 ± 6.1 |
| | Control | 2.3 – 31.7 | 10.7 | 12.7 ± 7.5 |
| Simpson diversity after 1 year | Reduced land use | 0.37 – 0.94 | 0.86 | 0.81 ± 0.14 |
| | Control | 0.31 – 0.95 | 0.83 | 0.79 ± 0.15 |
| Abundance after 3 years | Reduced land use | 648 – 4207 | 1644 | 1708 ± 726 |
| | Control | 172 – 2229 | 754 | 859 ± 577 |

The magnitude of treatment effects on abundance depended after one year (2021 sampling) and similarly after three years (2023 sampling) on past land use in the surrounding matrix and on how the reduced land-use plot had been mown in the previous year (Table 2, Fig. 3). In both years, effects of experimental land-use reduction became smaller when the matrix had in the previous years been mown more often, and statistical effect sizes in the averaged models were similar (2021: z=-3.383, p<0.001; 2023: z=-3.021, p=0.004). In turn, and again with comparable statistical effect sizes, treatment effects became larger on sites at more fertilized sites (2021: z=4.449, p<0.001; 2023: z=4.785, p<0.001). Regarding the specific type of mowing on the reduced land-use plot, log-response ratios after one year were larger when mowing was carried out with a greater cutting height (z=3.469, p<0.001) and after three years when the treatment was not mown at the same day as the matrix (z=2.830, p=0.007). Neither species richness, nor Shannon diversity nor Simpson diversity were significantly related to any variable in the averaged models. Cover of grassland and arable land and the size of the continuous grassland was not significant in any model.

**Fig. 3.**
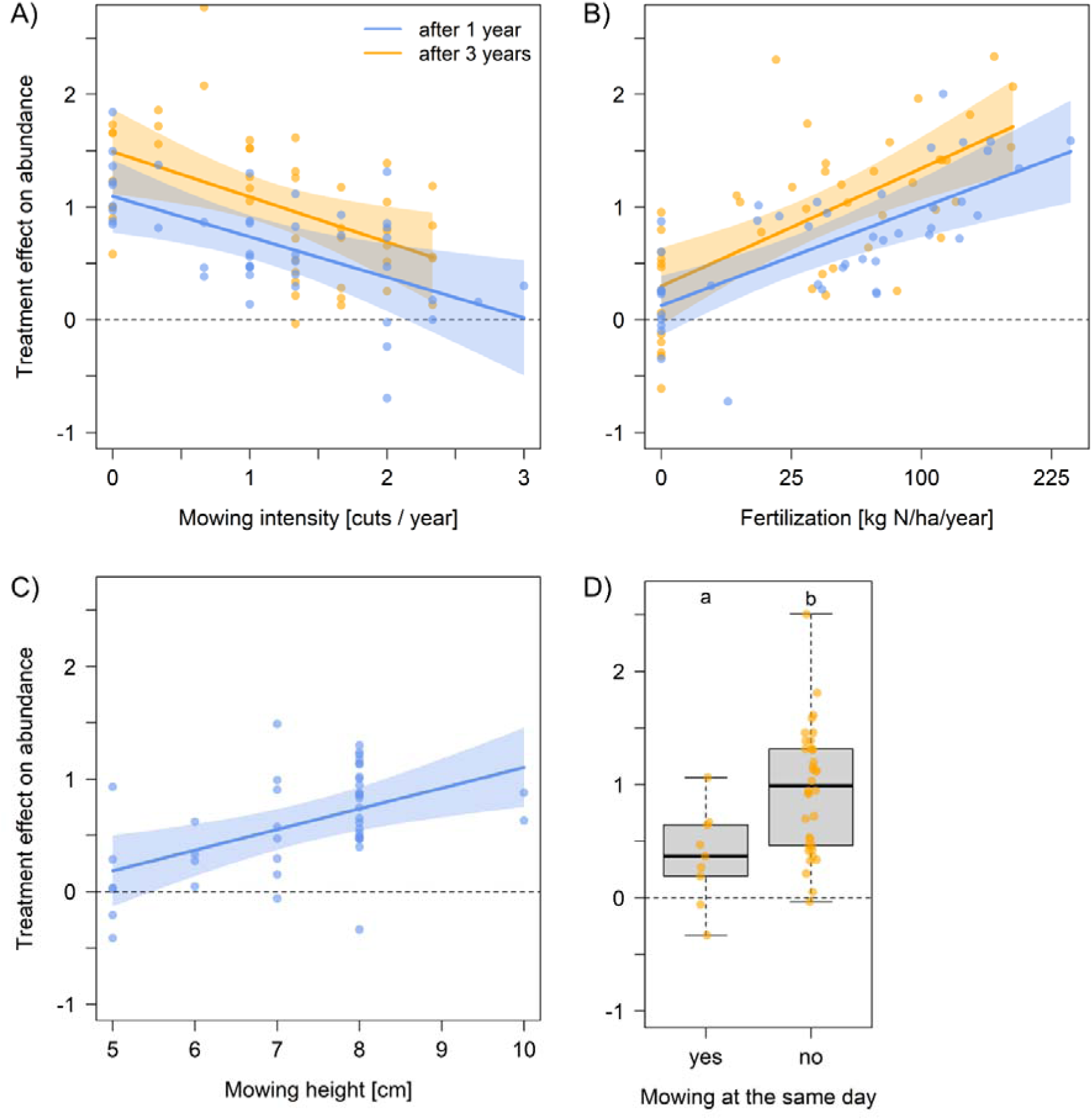
Treatment effects of experimental land-use reduction on arthropod abundance A) decreased in both sampling years when reduction plots were in a more frequently mown matrix. In turn, treatment effects became larger B) when the matrix had been fertilized more. One year after establishment of the experiment, C) treatment effects on abundance increased with mowing height of the reduced land-use plot; after three years, D) treatment effects were larger if reduced land-use and control plot had not been mown at the same day. Points are partial residuals of log-response ratios between treatment and control plots. Regression lines (95% CI as shaded polygons) indicate the predictions of averaged linear mixed-effects models. Dashed horizontal lines mark null. Mowing and fertilization are averaged over the respective three years before sampling. The x-axis in B) is on a square-root scale.

**Table 2.**
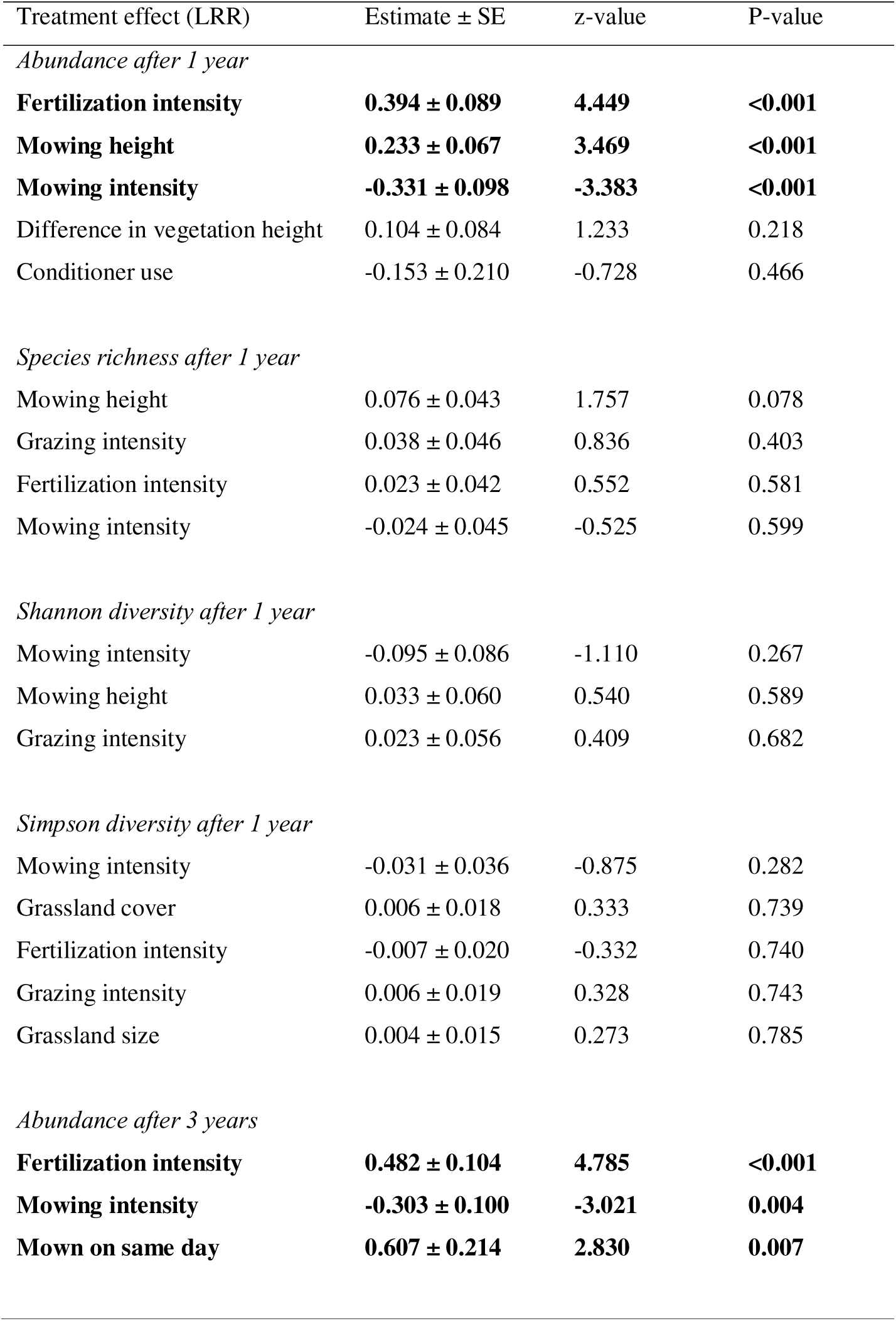
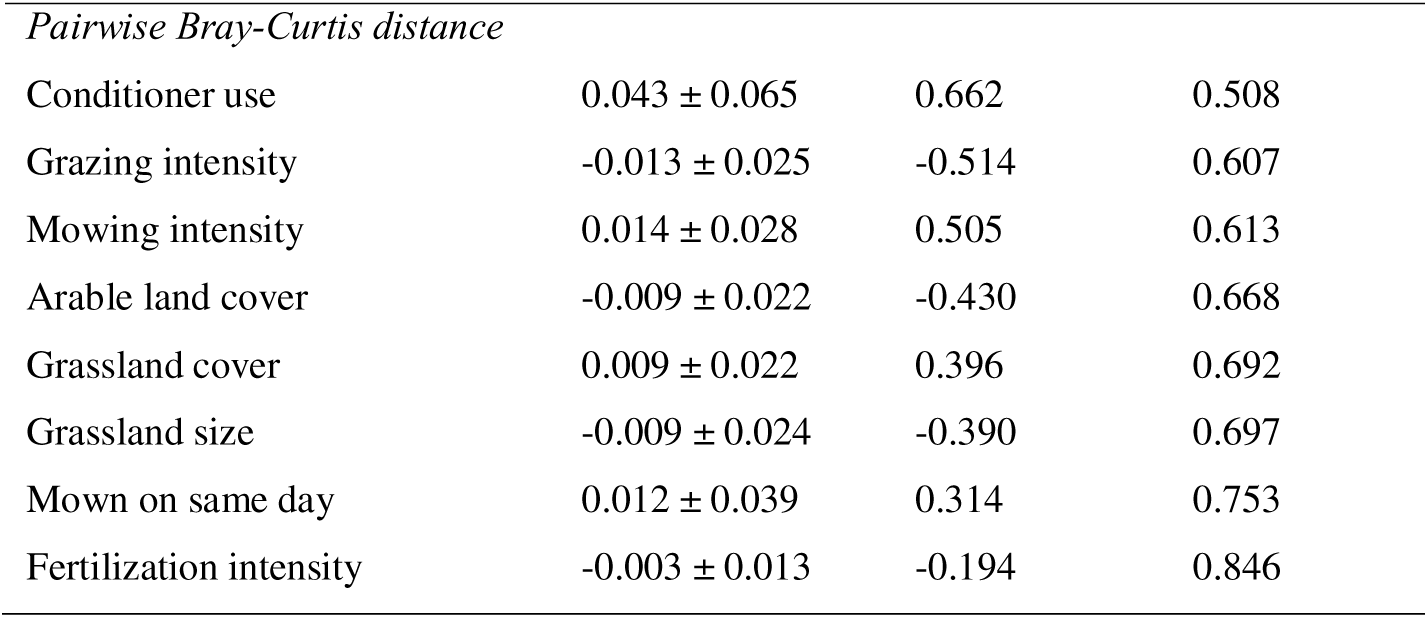
Results of the averaged (within _Δ_2 AICc units) linear models testing for the relationship between treatment effects of land-use reduction (quantified as log-response ratio, LRR) on invertebrate abundance, species richness, Shannon diversity, Simpson diversity, pairwise Bray-Curtis distance and explanatory variables. Reported are standardized model estimates ± standard errors (SE), z-values and P-values, with significant (at P < 0.05) relationships given in bold. Variables in each model are sorted by decreasing z-values.

| Treatment effect (LRR) | Estimate ± SE | z-value | P-value |
| --- | --- | --- | --- |
| *Abundance after 1 year* |  |  |  |
| **Fertilization intensity** | **0.394 ± 0.089** | **4.449** | **<0.001** |
| **Mowing height** | **0.233 ± 0.067** | **3.469** | **<0.001** |
| **Mowing intensity** | **-0.331 ± 0.098** | **-3.383** | **<0.001** |
| Difference in vegetation height | 0.104 ± 0.084 | 1.233 | 0.218 |
| Conditioner use | -0.153 ± 0.210 | -0.728 | 0.466 |
| *Species richness after 1 year* |  |  |  |
| Mowing height | 0.076 ± 0.043 | 1.757 | 0.078 |
| Grazing intensity | 0.038 ± 0.046 | 0.836 | 0.403 |
| Fertilization intensity | 0.023 ± 0.042 | 0.552 | 0.581 |
| Mowing intensity | -0.024 ± 0.045 | -0.525 | 0.599 |
| *Shannon diversity after 1 year* |  |  |  |
| Mowing intensity | -0.095 ± 0.086 | -1.110 | 0.267 |
| Mowing height | 0.033 ± 0.060 | 0.540 | 0.589 |
| Grazing intensity | 0.023 ± 0.056 | 0.409 | 0.682 |
| *Simpson diversity after 1 year* |  |  |  |
| Mowing intensity | -0.031 ± 0.036 | -0.875 | 0.282 |
| Grassland cover | 0.006 ± 0.018 | 0.333 | 0.739 |
| Fertilization intensity | -0.007 ± 0.020 | -0.332 | 0.740 |
| Grazing intensity | 0.006 ± 0.019 | 0.328 | 0.743 |
| Grassland size | 0.004 ± 0.015 | 0.273 | 0.785 |
| *Abundance after 3 years* |  |  |  |
| **Fertilization intensity** | **0.482 ± 0.104** | **4.785** | **<0.001** |
| **Mowing intensity** | **-0.303 ± 0.100** | **-3.021** | **0.004** |
| **Mown on same day** | **0.607 ± 0.214** | **2.830** | **0.007** |
| *Pairwise Bray-Curtis distance* |  |  |  |
| Conditioner use | 0.043 ± 0.065 | 0.662 | 0.508 |
| Grazing intensity | -0.013 ± 0.025 | -0.514 | 0.607 |
| Mowing intensity | 0.014 ± 0.028 | 0.505 | 0.613 |
| Arable land cover | -0.009 ± 0.022 | -0.430 | 0.668 |
| Grassland cover | 0.009 ± 0.022 | 0.396 | 0.692 |
| Grassland size | -0.009 ± 0.024 | -0.390 | 0.697 |
| Mown on same day | 0.012 ± 0.039 | 0.314 | 0.753 |
| Fertilization intensity | -0.003 ± 0.013 | -0.194 | 0.846 |

Likewise, difference in species composition per site (pairwise Bray-Curtis distances) were not related to any tested variable. Species composition in the NMDS ordination segregated according to land-use type, with grazed and mown sites being opposite (Fig. 4). Variation in NMDS scores was associated to the frequency of mowing (p=0.023; Appendix A: Table S3) and grazing intensity (p=0.031), which directly aligned along the first NMDS axis, and furthermore to grassland cover (p=0.029). Reduced land-use and control plots did not systematically differ in species composition (PERMANOVA, F=0.756, p=0.562) and multivariate dispersion (F=0.001, p=0.989).

**Fig. 4.**
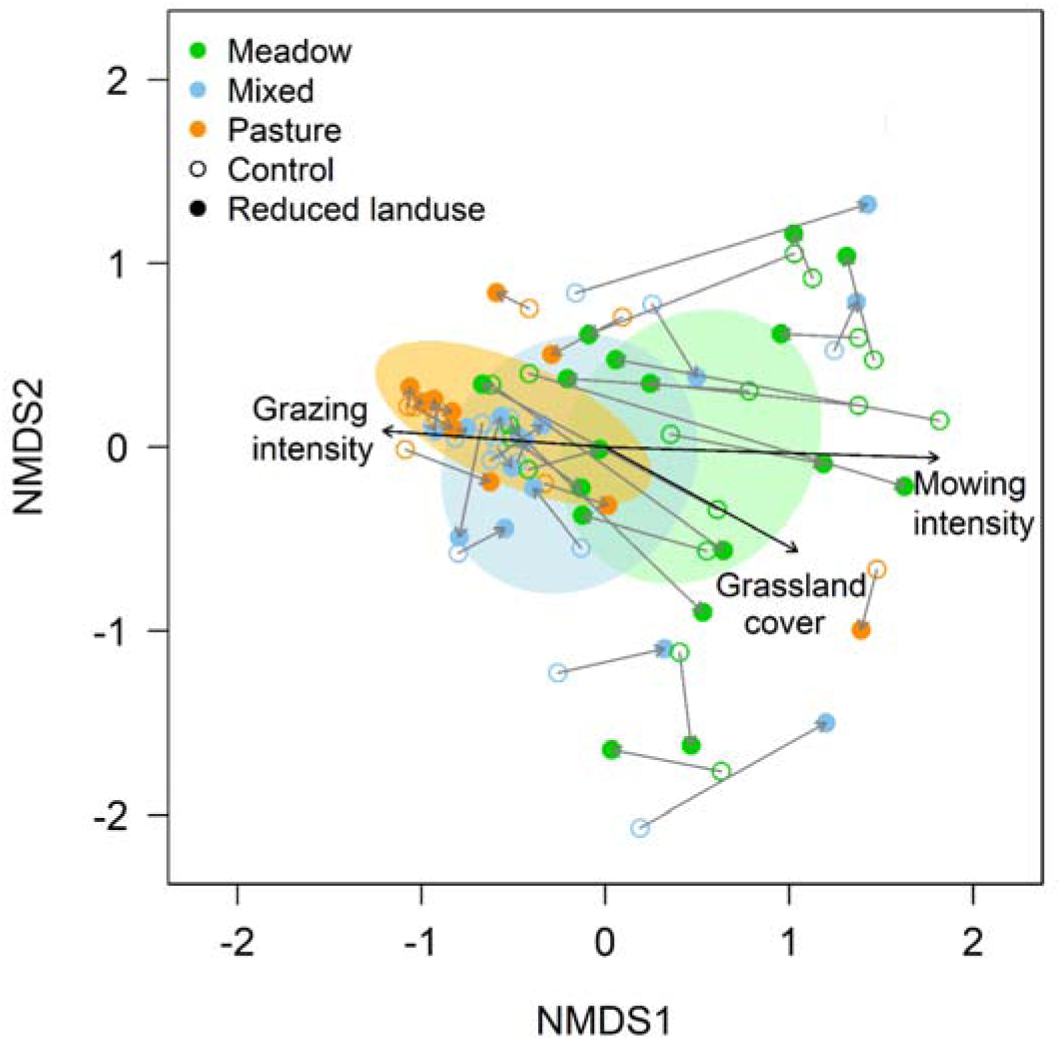
Species composition after one year (using meta-barcoding reads accounting for more than 0.2% of normalized reads per sample) did not systematically differ between reduced land-use and control plots but depended on the land use of the matrix, with meadows and pastures being opposite in the two-dimensional NMDS ordination (stress=0.205) and mixed sites being intermediate (land-use categories used for visualization only). Shaded polygons illustrate the 95% CI of ordination centroids per land-use category (meadow, mixed, pasture). Black arrows (obtained from a post-hoc permutation test, lengths of arrows proportional to strength of correlation) give the significant (at p<0.05) correlations between ordination scores and environmental variables (details in Appendix A: Table S3). Thin grey arrows connect control (open circles) and reduced land-use (filled circles) plots per site.

## Discussion

Experimental reduction of land-use intensity to a single late mowing increased invertebrate abundance – but not diversity – already after one year, and these positive effects strengthened over time. The magnitude of the effect of land-use reduction depended on the intensity and type of land use in the surrounding matrix, and also on the mowing practices of the reduction plot. This suggests that the success of grassland restoration for invertebrates varies across local contexts (Woodcock et al., 2021), similar to what has been observed for plants (Slodowicz et al., 2023). Importantly, sampling was conducted before the first land use on the sites and there was no relationship with differences in vegetation height between treatment and control. Hence, the experimental reduction plots in our study are not simple refugia and the results can be transferred to managed grasslands at large.

### Reduction in land use increases invertebrate abundance

The hypotheses that reduced land use increases both, invertebrate abundance and diversity, was only confirmed for abundance but not for diversity, irrespective of how rare or abundant species where weighted (species richness, Shannon diversity, Simpson diversity; sensu Hill, 1973). Finding consistently more individuals in grassland areas that had in the preceding year been used less intensively suggests that species already present at the sites benefit from reduced land use, rather than also being beneficial for additional species that are not yet part of the species pool in the surrounding matrix. This is also supported by the various composition analyses, as there was no systematic difference in composition between treatment and control. We admit that the land-use reduction experiment introduced mowing to grazed sites that were not mown in the past, even though grazing is frequently considered to be a type of land use that is less detrimental for invertebrates compared to mowing (Berger et al., 2024b; Chisté et al., 2016; Fartmann, 2024). Nevertheless, treatment plots on pastures did not become more similar to meadows, and meadows after being mown less did not converge towards pastures, which suggests a longer-lasting legacy of the preceding land use (Bürgi et al., 2017). A similar legacy is known for plants, which is why reduction and restoration may not immediately increase diversity or change composition (Humann-Guilleminot et al., 2023; Isbell et al., 2013). Likewise, plant composition in our experiment did not change in the first year (Andraczek et al., 2023). Consequently, invertebrate results were likely not driven by plants, as feeding resources for herbivorous insects were similar between treatment and control at each site, which may explain the absence of an effect on diversity and composition (sensu Schaffers et al., 2008).

The increase in abundance may be a direct response to reduced land use. Plausible explanations are, for example, that less frequent mowing killed fewer individuals (Berger et al., 2024b; Humbert et al., 2009) and that the absence of regular land use-associated disturbance resulted in more stable populations and higher reproductive output, increasing individual numbers per area. However, the lack of diversity differences might be explained by species that are losers of intensive land use and hence have not yet reestablish at the sites (Chisté et al., 2016; Mangels et al., 2017; Nickel & Achtziger, 2005). A taxonomic bias due to using meta-barcoding, is unlikely. Almost all (200 of 208) species of the taxa (Araneae, Auchenorrhyncha, Coleoptera, Heteroptera, Orthoptera) that have been monitored yearly on the control plots since 2008 (Seibold et al., 2019) had been collected and identified morphologically before from the sites. This indicates that our data are reliable and further supports that the species occurring in the reduced land-use treatment are those that have been present before, which then could increase in abundance.

The effect of land-use reduction became considerably larger in the second sampling, three years after the beginning of the experiment. Potentially, the yearly abundance increment in the reduction plots can add before eventually approaching the carrying capacity of the habitat (Chapman & Byron, 2018). Alternatively, and not mutually exclusive, the lasting legacy of intensive land use may gradually diminish, meaning that reducing land use will take several growing seasons to fully take effect (Humann-Guilleminot et al., 2023). However, we cannot fully rule out that the large increase in abundance in the second sampling was due to stochasticity, which is nevertheless unlikely, as large effect sizes for abundance were consistently found across all three study regions despite the local differences in climate or other site characteristics. Unfortunately, it was not possible to meta-barcode these specimens, so it remains unclear if after three years land-use reduction had changed diversity and composition.

### The magnitude of increase in abundance depends on land use

The increase of abundance in reduced land-use treatments was not constant but depended on land use in the surrounding matrix. In grasslands that were mown more often, reduction led to only few additional individuals, while in rarely mown or unmown grasslands reduced land-use plots had considerably more individuals. In contrast, for fertilization, increases were small in unfertilized sites and became larger when nutrient inputs were higher. This means that stopping fertilization had a larger effect on sites that previously experienced high fertilization levels compared with sites of low fertilization. Hence, the hypothesized greater treatment effect in less-intensive grasslands was not uniformly supported. Since mowing and fertilization were positively related (Spearman’s ρ = 0.671), due to the higher productivity and mowing potential in more fertilized grasslands, the differing relationships suggest different underlying mechanisms. As already elaborated on, every mowing event presents a high mortality risk to invertebrates in mown grasslands (Berger et al., 2024b; Humbert et al., 2009; Steidle et al., 2022), and highest abundance and diversity is often found in grasslands that are not mown at all (Frenzel & Fischer, 2022). When mowing is frequent, the continuous grassland matrix around the reduction treatment is depleted in individuals, which reduces source populations that can utilize the newly created reduction plots, and eventually establish populations therein. Thus, in frequently mown grasslands, reduction of land use may be less efficient, unless areas are connected to other unmown habitats or are larger than the 30 m x 30 m in our experiment, as the frequently mown matrix is likely a permanent sink habitat.

Compared to the likely direct impact of mowing, relationships with fertilization may be more indirect and mediated via plants. Increased nutrient supply raises the productivity of grasslands (Socher et al., 2012; Willems & van Nieuwstadt, 1996), which is at the expense of plant diversity, as a few nutrient demanding species become dominant and outcompete others via competition for light (Bobbink et al., 2010; Busch et al., 2019). If land use is now reduced and fertilization ceased, productivity will initially remain high and plant diversity and composition only start to recover once biomass and consequently nutrients are being removed (Isbell et al., 2013). Thus, at more intensively fertilized sites in our experiment, productivity in the reduction treatment may have been higher than in less fertilized sites, providing relatively more resources to invertebrates, which could hence reach higher densities. The magnitude of increase in abundance was not related to grazing, albeit large herbivores have profound effects on vegetation structure (Lundgren et al., 2024) and grazing can reduce invertebrate abundance and diversity (Helden et al., 2020; Kruess & Tscharntke, 2002). Nevertheless, grazing can maintain plant diversity (Eskelinen et al., 2022; Olff & Ritchie, 1998), and is often regarded as less negative for invertebrates compared to mowing (Fartmann, 2024), which may explain why invertebrate abundance in our data did not systematically vary with grazing intensity.

In addition to relationships with land-use intensity in the matrix, there was also an influence of how the experimental reduction plot was managed. Treatment effects on abundance were larger when the single late mowing was carried out at greater cutting height and when control and reduction plot were not mown simultaneously. The cutting height of mowing has previously been demonstrated to modulate insect mortality (Berger et al., 2024b; Hartlieb et al., 2024). Our results confirm that a greater height is less detrimental for invertebrates. When not the entire continuous grassland is mown at once at the same day, some of the area can serve as refuge and later be a source habitat from which individuals can spill-over into the mown part (Schwarz et al., 2023). Hence, a spatially and temporally non-uniform mowing regime has been recommended as beneficial for invertebrates in semi-natural grasslands (Fartmann, 2024; Johansen et al., 2019). In our experiments, a recolonization of the reduction plot from the surrounding matrix was probably more pronounced, when mowing was not on the same day, and relatively more individuals were then found in the subsequent year. Mechanistically, this result suggests that dispersal among habitat patches, the maintenance of meta-populations, and habitat connectivity are important for the restoration of insect populations in grassland (Nell et al., 2024; Panassiti et al., 2023; Rösch et al., 2013).

### Implications and outlook

Biodiversity in grassland has been declining (Hallmann et al., 2017; Jandt et al., 2022; Seibold et al., 2019). To halt and eventually reverse insect declines, reducing land-use intensity is important. However, as restoring species populations in grasslands may take a long time (Humann-Guilleminot et al., 2023), which is also confirmed by our data, it is pivotal to avoid degradation of grasslands that are not yet intensively used (Watson et al., 2016). Nevertheless in most European grasslands some management is required to prevent succession to scrub and woodland and to ensure the continued provisioning of specific services and commodities related to grasslands (e.g. fodder and milk production) (Shipley et al., 2024). This raises the question which land-use components should be prioritized in reduction programs, as mowing and grazing affect organisms differently (Schneider & Hering, 2024). While mowing at intermediate frequency can increase plant diversity by reducing the competitive ability of dominant plant species (Hautier et al., 2009), it is at the same time detrimental for invertebrates (Berger et al., 2024b; Humbert et al., 2009). In our experiment, the positive effect of extensification on invertebrate abundance was even more pronounced when the surrounding matrix was mown less frequently, and when mowing was down at greater cutting height and at different times. These results support that successful habitat management should be tailored according to the requirement of target organisms, because restoration may fail when key life histories are not addressed (see also Radinger et al., 2023; Thomas et al., 2009). Jointly and permanently maximizing restoration outcomes for plants and invertebrates may thus require a landscape approach with slightly different measures implemented in adjacent areas, also to increase heterogeneity of and connectivity among habitat patches (Allan et al., 2014; Grass et al., 2019).

Even though the number of individuals gained in reduction plots depended on the surrounding matrix, average effect sizes were positive, especially after three years. Hence, supporting invertebrate populations by lowering land-use intensity can be successful independent of specific local context. More extensively used grassland may furthermore be more resistant to climate change (Korell et al., 2024; Lyons et al., 2023) and increasing invertebrate abundance is expected to benefit consumer species such as insectivorous birds (Bowler et al., 2019) and to strengthen ecosystem functions (Weisser & Siemann, 2004). Nonetheless, reducing land use will have economic costs, and targeted subsidies may be necessary to implement land-use reduction at large scales. It will be particularly interesting to observe how outcomes of land-use intensity reduction will develop over time. Future research will clarify if invertebrate diversity in reduced land-use plots will eventually follow the direction of abundance once additional species had the chance to colonize and establish (Nickel & Achtziger, 2005). Extending sampling over more years will help to understand how restoration outcomes depend on drivers such as weather (Müller et al., 2024), and at which density individual numbers in reduced land-use plots saturate, which would further advance the scientific understanding of land-use reduction.

## CRediT authorship contribution statement

**Michael Staab**: Conceptualization; Data curation; Formal analysis; Investigation; Methodology; Project administration; Resources; Visualization; Writing - original draft; Writing - review & editing. **Alexander Keller**: Data curation; Formal analysis; Investigation; Methodology; Writing - review & editing. **Rafael Achury**: Conceptualization; Writing - review & editing. **Andrea Hilpert**: Data curation; Investigation; Methodology; Writing - review & editing. **Norbert Hölzel**: Conceptualization; Methodology; Writing - review & editing. **Daniel Prati**: Conceptualization; Project administration; Writing - review & editing. **Wolfgang W. Weisser**: Conceptualization; Funding acquisition; Project administration; Writing - review & editing. **Nico Blüthgen**: Conceptualization; Funding acquisition; Methodology; Project administration; Writing - review & editing.

## Declaration of competing interest

The authors declare that they have no known competing financial interests or personal relationships that could have appeared to influence the work reported in this paper.

### Ethics approval and consent to participate

Not applicable

### Consent for publication

Not applicable

### Data availability

This work is based on data elaborated by a project of the Biodiversity Exploratories program (DFG Priority Program 1374). The datasets are publicly available in the Biodiversity Exploratories Information System (http://doi.org/10.17616/R32P9Q) under the accession numbers 31767 (land use), 31966 (sampling meta data), 31967 (arthropod count data), and 31985 (meta-barcoding reads).

## Acknowledgments

We thank the managers of the three Exploratories, Julia Bass, Melissa Jüds, Anna K. Franke, Franca Marian, Max Müller, and all former managers for their work in maintaining the plot and project infrastructure; Victoria Grießmeier for giving support through the central office, Andreas Ostrowski for managing the central data base, and Markus Fischer, Eduard Linsenmair, Dominik Hessenmöller, Daniel Prati, Ingo Schöning, François Buscot, Ernst-Detlef Schulze, Wolfgang W. Weisser and the late Elisabeth Kalko for their role in setting up the Biodiversity Exploratories project. We are indebted to Jörg Hailer, Ralf Lauterbach, Uta Schumacher and Christin Schreiber for providing data and facilitating data collection, to Sarah Fritsch, Robin Fuchs, and Genevieve Walther for supporting arthropod collection. Sarah Sturm helped with preparing samples for meta-barcoding. We thank the administration of the Hainich national park, the UNESCO Biosphere Reserve Swabian Alb and the UNESCO Biosphere Reserve Schorfheide-Chorin as well as all land owners for the excellent collaboration. Field work permits were issued by the responsible state environmental offices of Baden-Württemberg, Thüringen, and Brandenburg

## Supplementary material

Supplementary material associated with this article can be found in the online version.

