## Appendix A for "Experimental reduction of land use increases invertebrate abundance but not diversity in grasslands"

**Contents**

**Table S1** Species numbers per order………………………………..…………..………………………………..….………..2

**Table S2** Pairwise contrasts for treatment effects………………………..………………………………..….………..3

**Table S3** Permutation test on NMDS axes scores…………….…………..………………………………..….………..4

**Fig. S1** Overview on data collection …………………………………………..……………………………………………..5

**Fig. S2** Correlations among treatment effects for different low-abundance filtering …………..…….6

**Fig. S3** Species accumulation curves ……………………………..……………..…………………………………..………7

**Fig. S4** Venn diagram for comparison with long-term monitoring……………………..…………………..…8

**Table S1.** Overview on species per order (or other higher-level taxonomic group) found with metabarcoding (applying a low-abundance filtering threshold of 0.2% normalized reads per sample).

| Order | Number of species |
| --- | --- |
| *Arachnida* |  |
| Araneae | 15 |
| Acariformes | 6 |
| *Collembola* |  |
| Entomobryomorpha | 19 |
| Symphypleona | 4 |
| *Myriapoda* |  |
| Diplopoda | 2 |
| Chilopoda | 1 |
| *Gastropoda* |  |
| Stylommatophora | 16 |
| *Insecta* |  |
| Coleoptera | 130 |
| Diptera | 124 |
| Hemiptera | 69 |
| Hymenoptera | 51 |
| Lepidoptera | 26 |
| Orthoptera | 9 |
| Strepsiptera | 1 |
| Thysanoptera | 1 |
| *Malacostraca* |  |
| Isopoda | 5 |
| Total | 479 |

**Table S2.** Pairwise contrasts for treatment effects (log-response ratios) between invertebrate abundance, species richness, Shannon diversity and Simpson diversity for the sampling in 2021 (after one year). P-values are Bonferroni-Holm-corrected for multiple comparisons. Significant contrasts (p<0.05) are printed in bold.

| Contrast | Estimate ± SE | t-ratio | p-value |
| --- | --- | --- | --- |
| **Abundance – Species richness** | **0.382 ± 0.070** | **5.436** | **<0.001** |
| **Abundance – Shannon diversity** | **0.357 ± 0.070** | **5.078** | **<0.001** |
| **Abundance – Simpson diversity** | **0.388 ± 0.070** | **5.516** | **<0.001** |
| Species richness – Shannon diversity | -0.025 ± 0.071 | -0.356 | 1.000 |
| Species richness – Simpson diversity | 0.006 ± 0.071 | 0.080 | 1.000 |
| Shannon diversity – Simpson diversity | 0.031 ± 0.071 | 0.435 | 1.000 |

**Table S3.** Results of the post-hoc permutation test (n=10,000 permutations) relating the scores of the first two NMDS axes (see Fig. 4) to environmental variables. Significant (at p<0.05) correlations are given in bold. Variables are sorted by decreasing R².

| Variable | NMDS1 | NMDS2 | p-value | R² |
| --- | --- | --- | --- | --- |
| **Mowing intensity** | **0.999** | **-0.033** | **0.023** | **0.265** |
| **Grassland cover** | **0.881** | **-0.474** | **0.029** | **0.216** |
| **Grazing intensity** | **-0.997** | **0.073** | **0.031** | **0.176** |
| Vegetation height | 0.853 | 0.521 | 0.253 | 0.163 |
| Grassland size | -0.599 | -0.800 | 0.276 | 0.162 |
| Fertilization intensity | 0.608 | 0.794 | 0.359 | 0.096 |
| Arable land cover | -0.921 | -0.390 | 0.794 | 0.093 |
| Treatment | 0.028 | -0.011 | 0.885 | 0.008 |

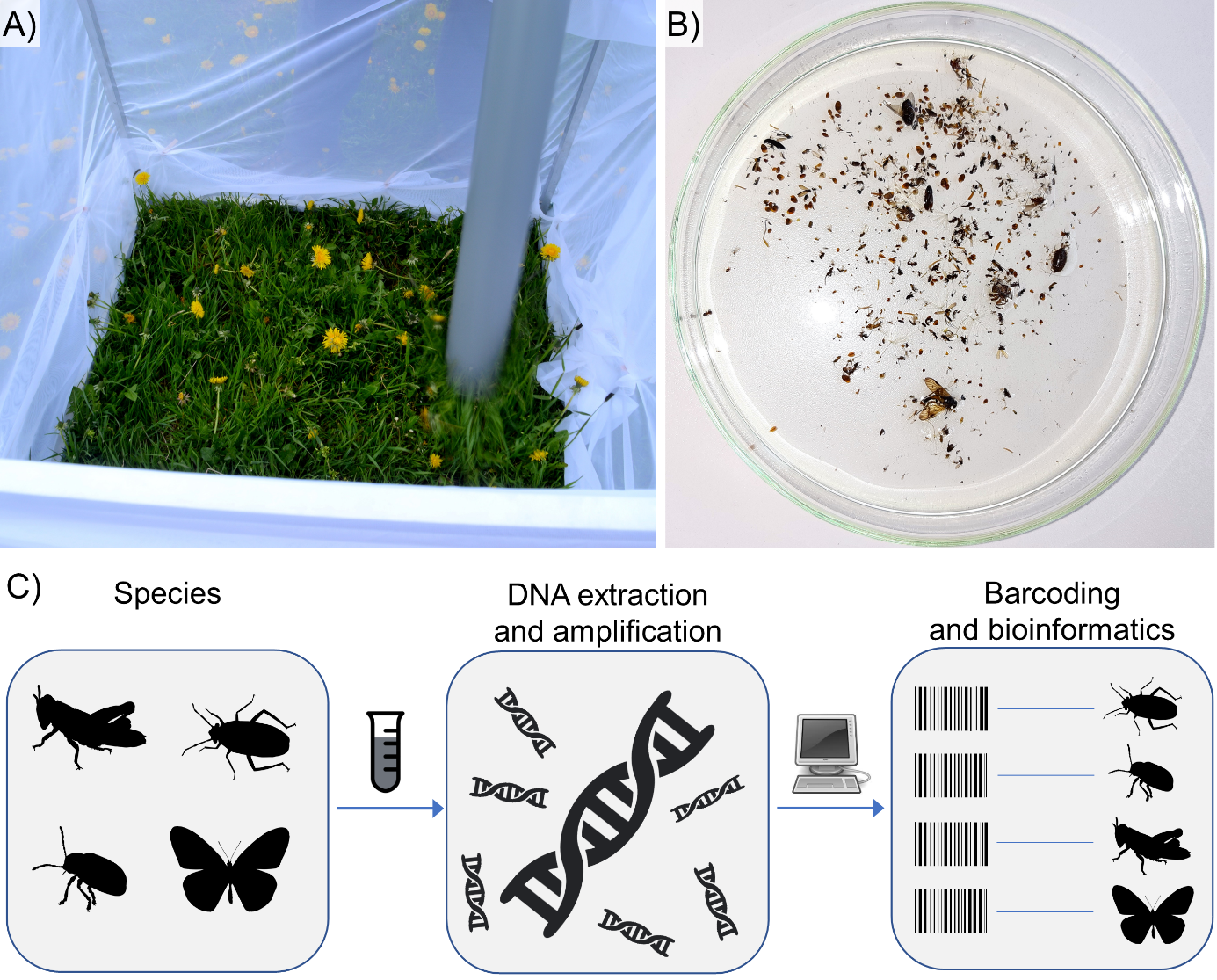

**Fig. S1.** Overview on data collection. A) Arthropods (and other invertebrates) were quantitatively collected by suction-sampling. Per plot, two 1 m² samples were taken. B) Bulk specimens were segregated from debris, sorted and counted. C) Species identification was accomplished by meta-barcoding. Pictures in a) and b) by Michael Staab; arthropod silhouettes in C) from phylopic.org, freely available under a CC0 1.0 license; icons in C) from openclipart.org, freely available under a CC0 1.0 license.

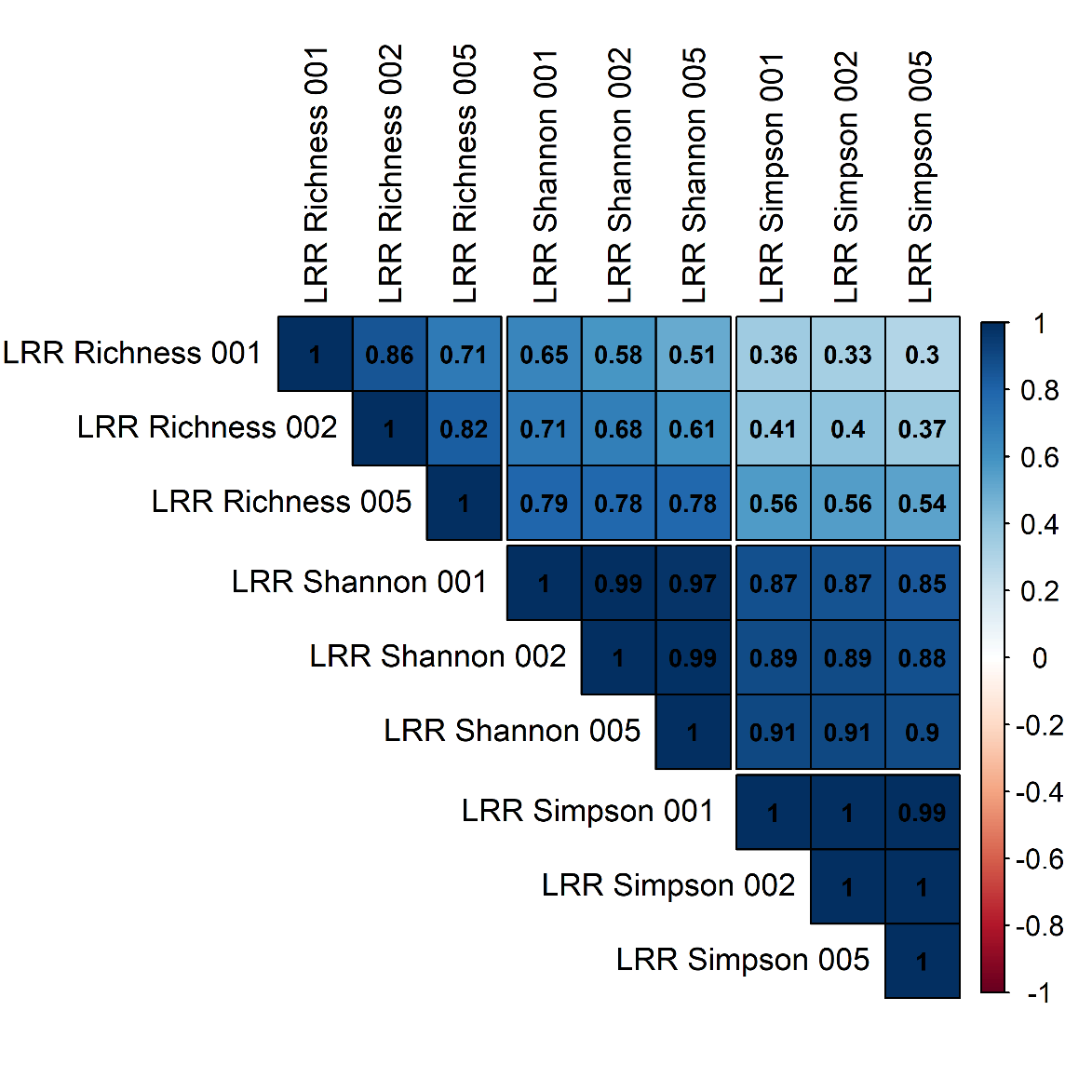

**Fig. S2.** Correlations (Spearman’s *ρ*) among log-response ratios (LRR) of species richness, exponential Shannon diversity and Simpson diversity, calculated for different low-abundance filtering omitting reads contributing to either less than 0.1%, 0.2% or 0.5% of normalized reads per sample. Correlations of differently-filtered diversity variables for the same type of diversity were consistently high, indicating that the results are not influenced by rare sequences.

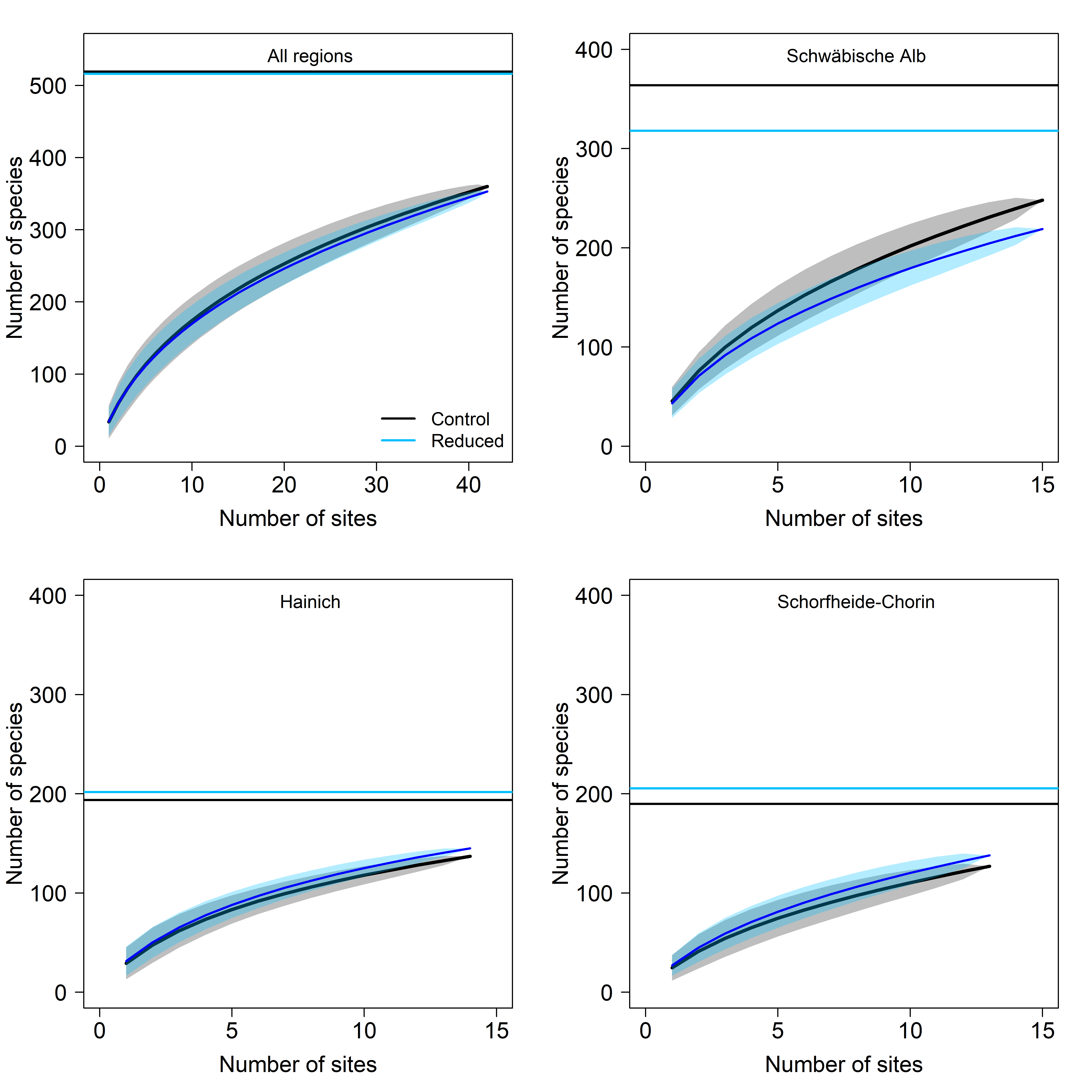

**Fig. S3.** Sample-based species accumulation curves (based on n=10,000 permutations, shaded polygons indicate 95% CI) of control (in black) and reduced land-use plot (i.e. experimental treatment; in blue) for all regions combined and separated per region (using meta-barcoding reads accounting for more than 0.2% of normalized reads per sample). Horizontal lines indicate expected number of species based on first-order jackknife estimators. Sampling efficiency was always similar (ranging between 68% and 72%), did not differ between control and treatment, and was congruent across regions.

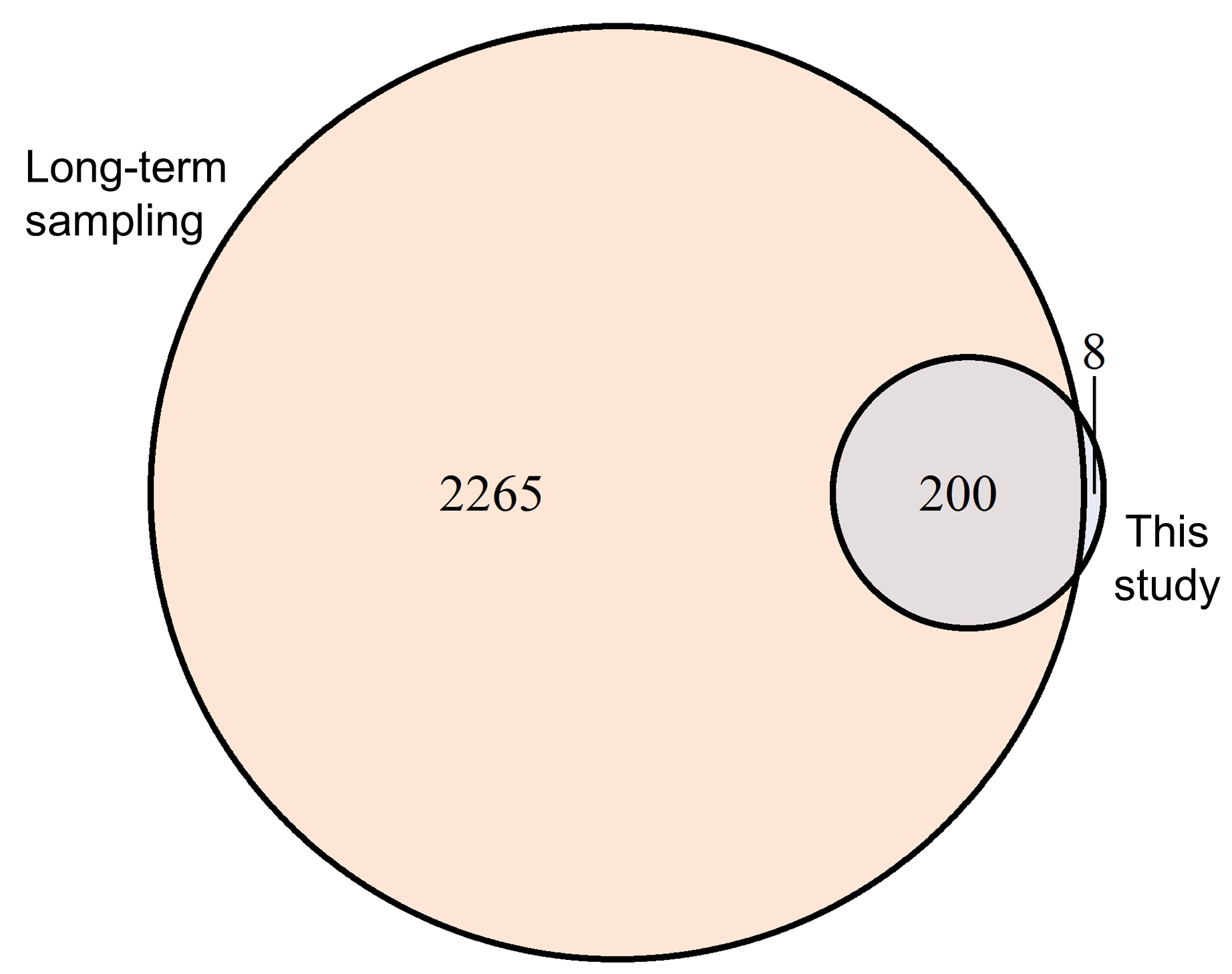

**Fig. S4.** Venn diagram showing that the majority of species (96%) from Araneae, Coleoptera, Hemiptera (comprising Auchenorrhyncha and Heteroptera) and Orthoptera (i.e. the higher-level taxa regularly monitored on the control plots) recorded by meta-barcoding in this study, had been known from morphological identifications in the long-term monitoring before. This gives credibility to the use of meta-barcoding, especially because sampling by biocenometer was previously not used on the sites. Data shown are based on sequences that account for more than 0.2% of normalized reads per sample.
